# Engineered sex distortion in the global agricultural pest *Ceratitis capitata*

**DOI:** 10.1101/2020.08.07.240226

**Authors:** Angela Meccariello, Flavia Krsticevic, Rita Colonna, Giuseppe Del Corsano, Barbara Fasulo, Philippos Aris Papathanos, Nikolai Windbichler

**Affiliations:** Department of Life Sciences, Imperial College London, Sir Alexander Fleming Building, South Kensington Campus, London, United Kingdom; Department of Entomology, Robert H. Smith Faculty of Agriculture, Food and Environment, Hebrew University of Jerusalem, Rehovot, Israel

## Abstract

Genetic sex ratio distorters have potential for the area-wide control of harmful insect populations. Endonucleases targeting the X-chromosome and whose activity is restricted to male gametogenesis have recently been pioneered as a means to engineer such traits. Here we enabled endogenous CRISPR/Cas9 and CRISPR/Cas12a activity during spermatogenesis of the Mediterranean fruit fly *Ceratitis capitata*, a worldwide agricultural pest of extensive economic significance. In the absence of a chromosome-level assembly, we analysed long and short-read genome sequencing data from males and females to identify two clusters of abundant and X-chromosome specific sequence repeats. When targeted by gRNAs in conjunction with Cas9 they yielded a significant and consistent distortion of the sex ratio in independent transgenic strains and a combination of distorters induced a strong bias towards males (~80%). Our results demonstrate the design of sex distorters in a non-model organism and suggest that strains with characteristics suitable for field application could be developed for a range of medically or agriculturally relevant insect species.

## Introduction

Effecting a substantial bias in the reproductive sex ratio of a population towards males has long been recognized as a potentially powerful means for genetic control^1^. While naturally occurring sex distortion traits have been of longstanding interest to evolutionary biologists^2^, the advent of powerful genome-editing tools and synthetic biology now allows for the generation of artificial sex distortion traits designed to realize this untapped potential. Sexually reproducing insect species that vector disease or disrupt agricultural production are considered the prime targets for such a genetic control approach that, in its various flavours, is predicted to be more efficacious than the mass release of sterile males^3^, the current gold standard in genetic pest control^4^. Several molecular mechanisms could be engineered to bring about a genetic male bias, which is preferred as females are generally the sex causing damage or transmitting disease. X-shredding has been pioneered in the mosquito *Anopheles gambiae* due to the fortuitous discovery of an X-linked target gene cluster and a suitable endonuclease^5,6^. X-shredding interferes with the transmission of the X-chromosome by inducing multiple DNA double strand breaks during male meiosis. More recently, the advent of CRISPR/Cas9 enabled the design of RNA-guided X-shredders that have now successfully been demonstrated in *Anopheles gambiae^7^* and *Drosophila melanogaster^8^*. As an alternative approach the male-determining genes or M-factors that have recently been identified in several insect species could also possibly be used to engineer effective sex conversion traits^9–13^.

Artificial sex distorters could be applied in a variety of ways, some of which have now been demonstrated in the laboratory. Autosomal distorter traits have been shown to eliminate caged mosquito populations^6^ and could, possibly in the form of repressible or inducible transgenes, be developed as a next generation inundative genetic control strategy for insects that are currently controlled by the sterile insect technique (SIT). Both X-shredders and M-factors could be mobilized via gene drive for a powerful inoculative type of control as has been recently demonstrated in *Anopheles gambiae^14^*. Similarly, when linked to the Y-chromosome, X-shredders could constitute a driving Y-chromosome that would favour its transmission over the X-chromosome and the resulting decline in female numbers is predicted to lead to the eventual collapse of the population^15^. This strategy was one of the first proposed but may be challenging to engineer due to transcriptional silencing of the Y chromosome during male meiosis^16^. Finally, post-zygotic genetic sex distorters that act in and are transmitted via the male germline could also be linked to the Y-chromosome and would obtain a level of persistence in the population. Such post-zygotic sex ratio distorters have recently been demonstrated in *Drosophila melanogaster* via the targeting of X-linked haplolethal genes during male meiosis^8^.

The Mediterranean fruit fly (medfly, *Ceratitis capitata*) is one of the most destructive global agricultural pests due to its wide host range of close to 300 different crop plants^17^. SIT is widely applied as the primary biological method to control medfly populations as a component of area-wide integrated pest management^18^. Nevertheless, the global economic costs due to crop damage, export controls due to quarantine restrictions, and control and prevention of medfly infestation are well over US$ 1 billion annually^19^ making the medfly an excellent candidate for further development of novel genetic control strategies.

We have recently identified the Y-linked M-factor as one possible avenue to engineer sex-distortion traits in the medfly^12^. Here we sought to explore the possibility of engineering CRISPR-based sex-distortion traits following the X-shredding paradigm, which operates during male meiosis to increase the transmission of Y-bearing gametes and does not entail the manipulation of the endogenous sex determination pathway. No chromosome-level assembly is currently available for *C. capitata* and although CRISPR/Cas9 ribonucleoprotein complexes have been successfully used to induce heritable genetic changes in the medfly^20^, there are currently no available tools for the endogenous expression of CRISPR. The establishment and demonstration of the X-shredding mechanism in the medfly would thus not only add further evidence that this mechanism is broadly applicable but also that such traits can be established in other organisms of great medical or economic importance, for which a full suite of genomic resources and a powerful set of genome editing tools are currently lacking.

## Results

### Identification of targetable sequence repeats on the medfly X-chromosome

We have previously described the redkmer pipeline which identifies putative X-linked repeat sequences that are absent from other chromosomes using whole genome sequencing reads as its primary input^21^. Here we used available medfly WGS data^12^ consisting of ~1.9M PacBio reads from male medflies of the Fam18 strain and approximately 4.9×10^8^ Illumina reads from each sex to identify X-linked sequences which could be targeted by CRISPR/Cas9^22^ or CRISPR/Cas12a^23^ in the male germline. Starting from a set of 5.3×10^8^ kmers of 25 nucleotides, redkmer selected 5×10^4^ kmers representing the top 0.05% most abundant and X-chromosome specific kmers (Figure 1A). We applied further rounds of selection to this initial set. Targetability by either Cas9 or Cas12a was predicted by FlashFry^24^ after which we applied criteria that ensure that selected kmers are not only highly abundant in both Illumina and PacBio libraries, but also represented on a maximum number of independent PacBio reads (Figure 1B,C & D). This was done to select against spurious repeats arising from sequencing artefacts and for those where individual repeat units are sufficiently spaced for independent targeting by Cas9 or Cas12a. From the resulting top 25 kmers we manually selected two Cas9 and two Cas12a kmers for the experimental validation steps (Supplementary Table S1, Figure 1B).

**Figure 1.**
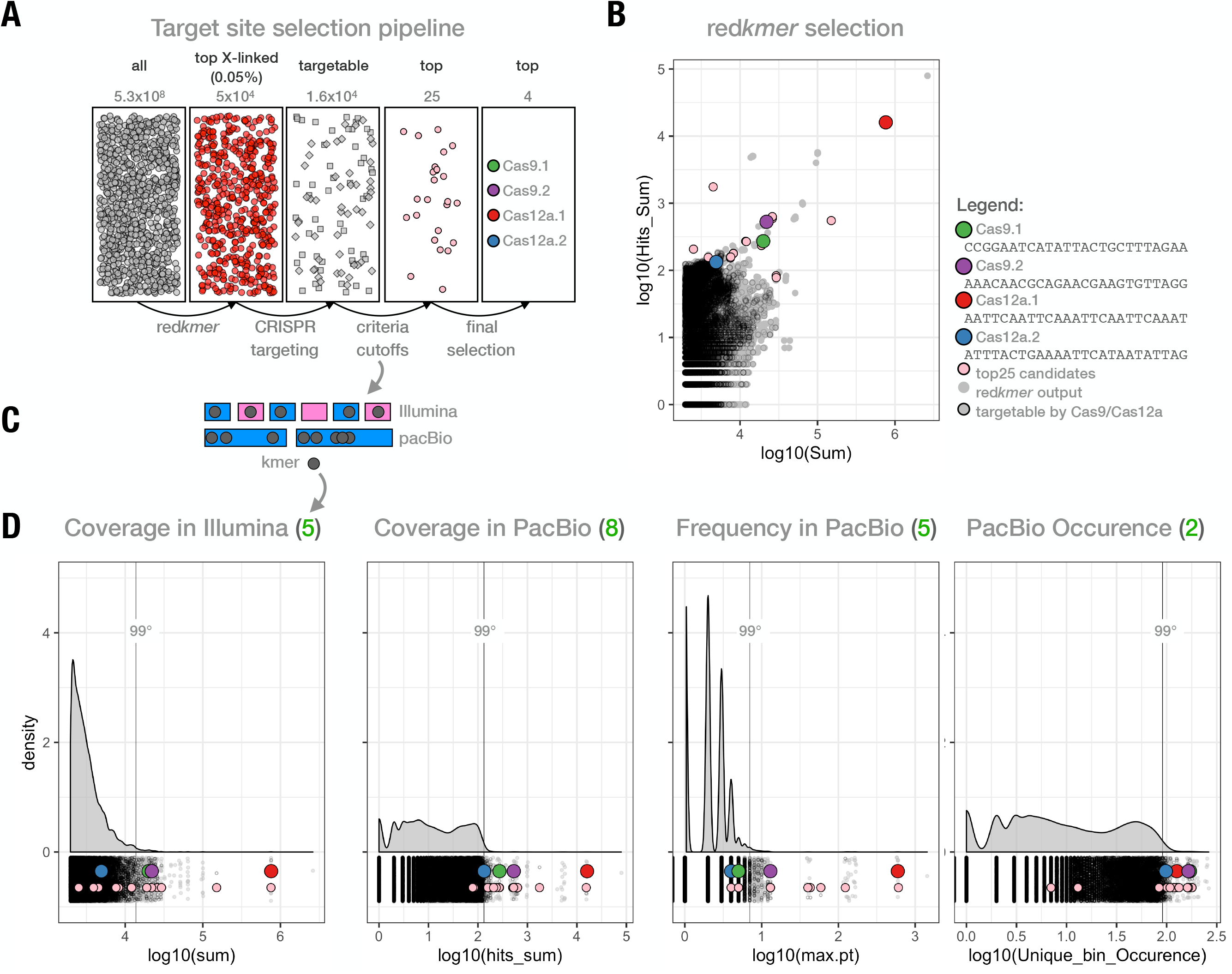
X-shredding target site selection. **A)** Overview of kmer selection pipeline and steps involved in the final selection. **B)** Plot showing coverage in Illumina (sum) and PacBio (Hits_Sum) datasets of all redkmer output candidate X-kmers. The top 25 selected and targetable kmers resulting from the manual downstream selection are shown in pink and those experimentally validated shown along with their sequence. **C)** Schematic representation of selection criteria used and how they relate to the Illumina (male and female) and PacBio (male only) datasets shown here for an example kmer (grey). **D)** Distribution of redkmer selected candidate X-kmers (in grey) for the selection criteria including the top25 and tested kmers. The green number in parenthesis next to the criterion label (top) shows the respective value of the example kmer shown in C and its interaction with the example Illumina and PacBio libraries.

### Activity of Cas9 and Cas12a activity during medfly spermatogenesis

To evaluate CRISPR function in the male germline of the medfly, we generated piggyBac transformation constructs for random integration in which Cas9 or Cas12a coding sequences were placed under the transcription control of the previously characterized *C. capitata* beta2-tubulin promoter^25^. In both the *An. gambiae* and *D. melanogaster* this promoter drives expression of genes during the primary spermatocyte stage at the onset of meiosis I and endonuclease activity at this stage has been demonstrated to cause shredding of the X-chromosome and to interfere with its transmission to progeny^7,8^. For the expression of the gRNA we identified a putative U6 gene using BLAST of the *Drosophila* U6snRNA sequence (GenBank accession no. NR002083) against the medfly genome (taxid:7213). A candidate medlfy promoter element was then obtained by gene synthesis. In addition to two constructs each harbouring a gRNA targeting one of the two top Cas9 candidate repeat sequences (Cas9.1 and Cas9.2), we also generated constructs with a previously-described gRNA^20^ targeting the autosomal *white eye* (Cas9.w) gene for evaluating CRISPR activity (Figure 2A). Two Cas12a constructs were generated, a construct targeting only the *white eye* gene (Cas12a.w), and a multi-gRNA construct (Cas12a.m) containing both gRNAs for the top X-shredding targets in addition to the *white eye* gRNA (Figure 2A). We generated multiple independent transgenic medfly strains using piggyBac mediated random integration. We performed inverse PCR to characterize insertion sites (Supplementary Figure 1, Supplementary Table S3). A total of 7 and 4 independent strains were established for the constructs targeting Cas9 kmer 1 and Cas9 kmer 2, respectively, and single strains for all other constructs. To assess Cas9 and Cas12a activity against the *white eye* marker gene we crossed transgenic sons or daughters of Cas9.w, Cas12a.w or Cas12a.m females to the w2Δ strain which carries a deletion spanning exon 2 of the *white eye* gene^26^. Mutatgenesis of the *C. capitata white eye* by transient CRISPR/Cas9 targeting of a conserved domain within exon 3 has been previously shown to result in a high rate of individuals lacking eye pigment^20^. We compared the activity of the two endonuclease platforms by counting the fraction of offspring with white eyes compared to the female control in which we did not expect endonuclease expression (Figure 2). In the progeny of transgenic males, we observed on average 98% white-eyed individuals for Cas9.w strain and <25% for strain Cas12a.w. (Figure 2B,C). We scored individuals as mutant only when pigment loss was complete (Figure 2C), but not when we observed mosaic or orange eyes which were infrequently observed in the progeny of Cas12a.w males. We detected no white-eyed individuals in the cross using the multiplexed strain Cas12a.m and also in a control where we crossed wild type individuals to the w2Δ strain (Figure 2). We found evidence for limited Cas9 activity in the Cas9.w female control cross with up to 0.34% mutant progeny detected in one family. This suggests that some low level of leaky somatic expression of the beta2-tubulin driven transgene can occur in females. Genotyping and sequencing of mutant *white eye* alleles from strain Cas9.w showed a spectrum of indels at the expected target site within exon 3 (Figure 2D). We concluded that, using the beta2-tubulin promoter in combination with Cas9, we were able to achieve high rates of activity during medfly spermatogenesis, although some level of ectopic expression in females could not be excluded.

**Figure 2.**
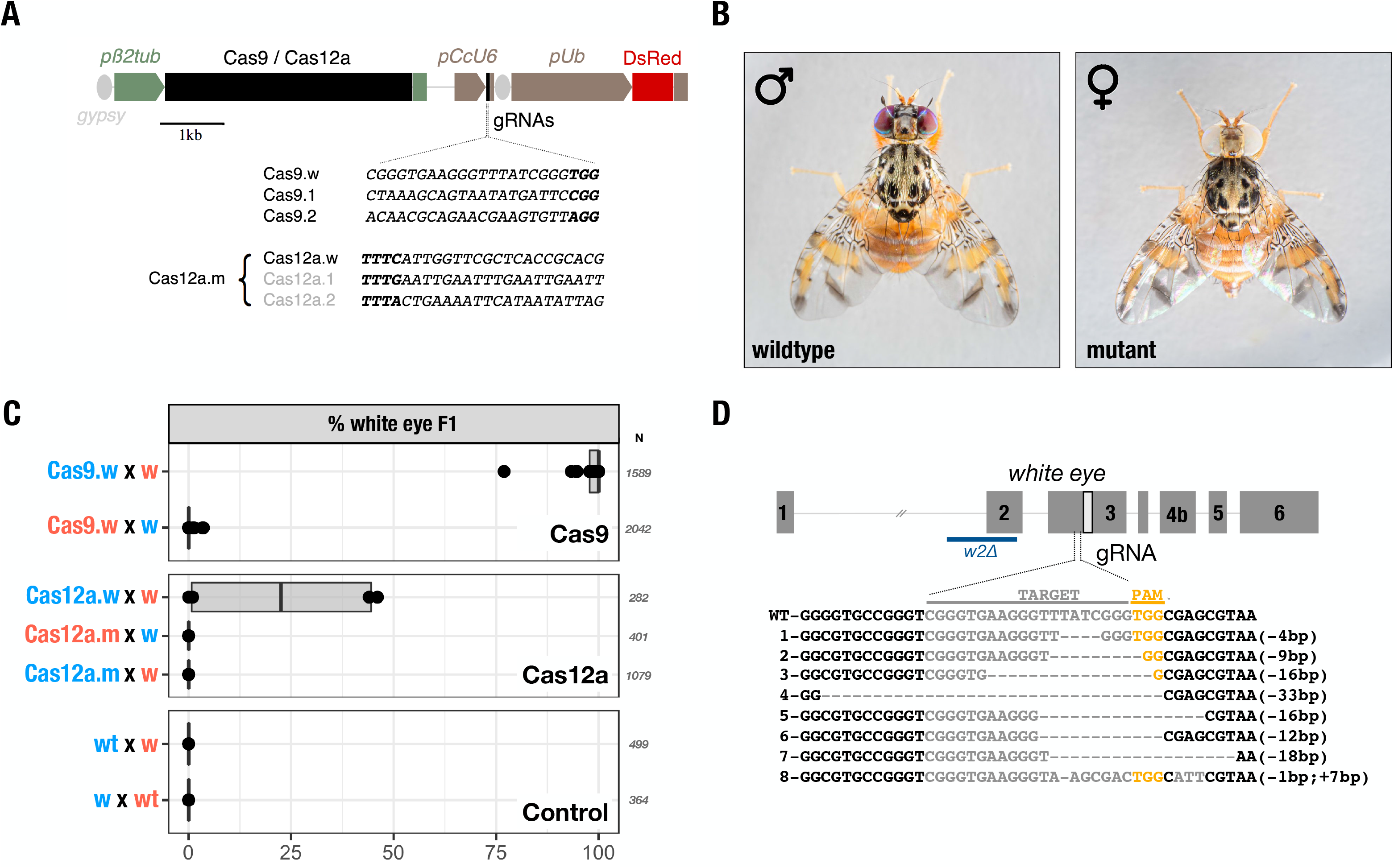
CRISPR/Cas constructs and activity against white eye in the male germline. **A)** Schematic representation of the transformation constructs. Cas9 or Cas12a coding sequences are under the transcriptional control of the male germline-specific *β2 tubulin promoter (pβ2tub)*. The gRNA target sequences are under the control of the endogenous *U6 Pol III promoter (pCcU6)* and the constitutively expressed DsRed as marker of transgenesis is under the control of the *Ubiquitin* promoter *(pUb)*. *Gypsy* insulator sequences (*gypsy*) are indicated adjacent to *pβ2tub* and *pUb*. **B)** Wildtype red-eye phenotype of an adult male medfly (left) and white-eye phenotype of a female medfly (right). **C)** The percentage of white-eyed flies obtained by individually crossing hemizygous males expressing Cas9 and the gRNA targeting the *white eye* gene (Cas9.w) with white-eyed w2Δ/w2Δ females (w) is shown in the top panel. As a control, hemizygous females (Cas9.w) were individually crossed with 10 white-eyed w2Δ/w2Δ males (w). The middle panel, shows the percentage of white-eyed flies obtained by individually crossing hemizygous males expressing Cas12a and the gRNA targeting the *white eye* gene (Cas12a.w) with white-eyed w2Δ/w2Δ females (w); by crossing hemizygous males for Cas12a.m with white-eyed w2Δ/w2Δ females (w) or by crossing hemizygous Cas12a.m females with w2Δ/w2Δ white-eyed males (w). The bottom panel shows control crosses between wildtype males with white-eyed females from the w2Δ strain (wt x w) and white-eyed males from the w2Δ strain crossed with wildtype females (w x wt). The numbers indicate the total number of individuals scored (N). **D)** Diagram representation of the autosomal *white eye* gene of *Ceratitis capitata*, spanning six exons, and the sequence of the gRNA targeting exon 3 (PAM is shown in yellow). Below, the wildtype sequence is aligned to sequences obtained from F1 white-eyed flies, showing the indels induced by CRISPR/Cas9. The deletion spanning exon 2 of the w2Δ *white eye* mutant strain is indicated.

### Characterization of candidate X-shredding strains

Transgenic individuals carrying gRNAs targeting Cas9 kmer1 (Cas9.1 strains a to g), those targeting Cas9 kmer2 (Cas9.2 strains a to d) as well as the Cas12a.m strain were crossed to the wild types to determine the sex-ratio of their offspring (Figure 3A). We expected to observe an effect in males due to the strong expression of the endonuclease during spermatogenesis and the reciprocal cross, due to the comparatively low levels of *β2* tubulin activity in females, was used as the control. Figure 3A summarizes the results of these experiments performed over four consecutive generations of batch crosses. We found that all Cas9 strains displayed a significant bias towards males in the progeny of transgenic fathers, but not in the progeny of transgenic mothers. An exception was strain Cas9.2a in which a significant deviation from the expected 50% sex ratio was also observed in the progeny of female transgenics, albeit less pronounced (55% males) compared to the male transgenic cross (66% males). Since this was not seen in the other Cas9.2 strains carrying the same construct we attributed this to a position/integration effect which may or may not be causally linked to the action of Cas9. No detectable effect on the sex-ratio was observed for strain Cas12a.m. Cas9.1 strains showed a male-biased sex-ratio of 54% on average whereas all Cas9.2 strains scored a highly-significant and more substantial sex bias of on average 66% male progeny with the best performing strain Cas9.2b reaching a bias of 68% towards males. We measured egg-to-adult survival rates for all Cas9 strains to determine whether targeting these two clusters had a significant postzygotic effect on survival. We found highly variable rates of survival across experimental groups including also the two wild type controls. To detect effects of X-shredding rather than strain-specific or insertion related effects we combined data from all Cas9.1 and from all Cas9.2 strains. Overall, a significant decrease in egg-to-adult survival was observed for Cas9.1 but not Cas9.2 although the latter set of strains shows a consistently more biased sex ratio towards males (Figure 3B).

**Figure 3.**
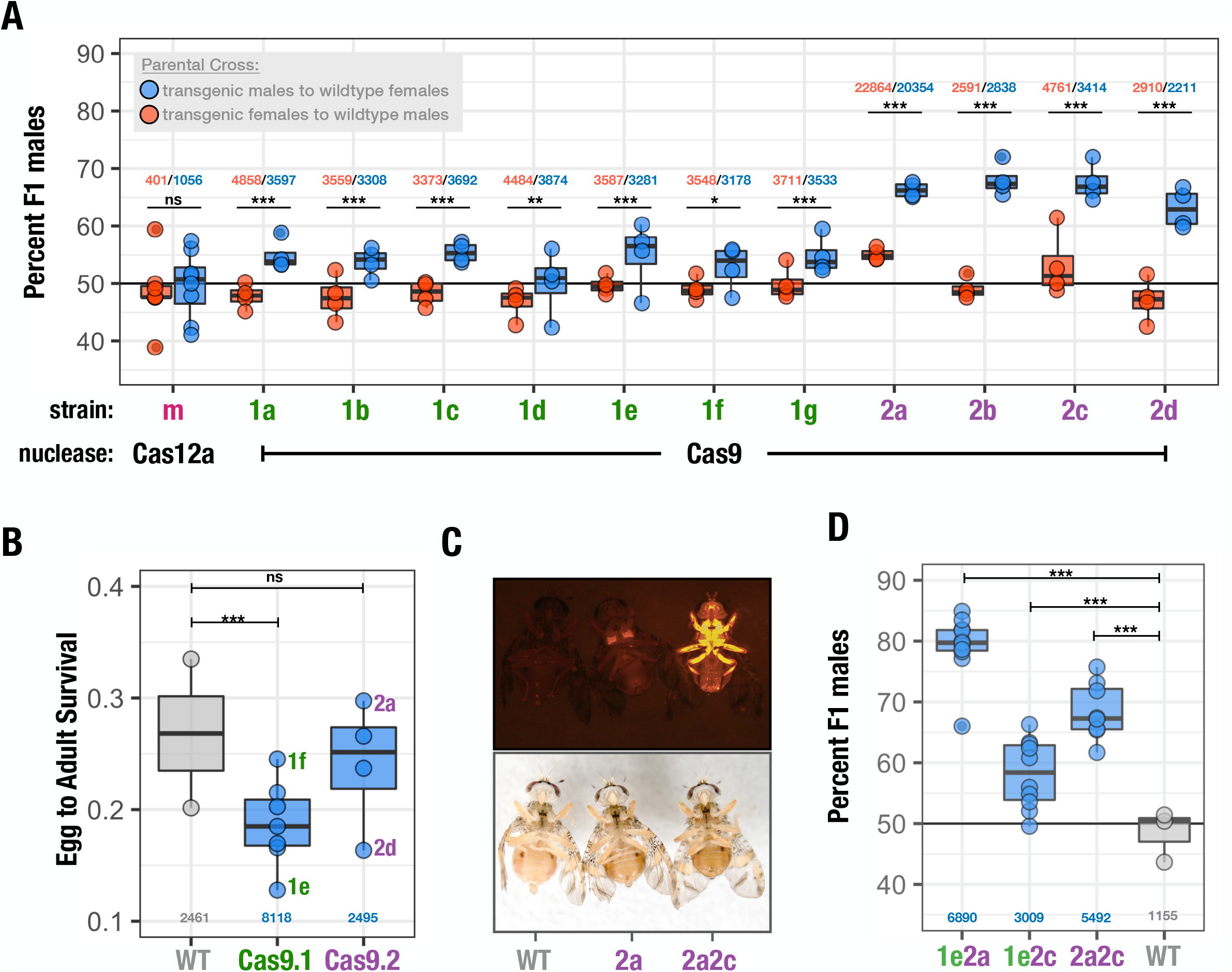
Analysis of the reproductive sex-ratio in Cas9 and Cas12a transgenic strains. **A)** Percentage of adult males in the progeny of transgenic males (blue) or females (red) from each transgenic strain crossed to the wild-type. Each strain was assayed over for four consecutive generations. **B)** Egg-to-adult survival rate of the progeny of transgenic males crossed to wildtype females of the Cas9.1 and Cas9.2 transgenic strains compared to the wildtype. **C)** A wildtype male (WT), a Cas9.2a hemizygous transgenic male carrying a single autosomal transgene (2a) and a Cas9.2a/Cas9.2c transhemizygote male carrying two autosomal transgenes (2a2c) observed using the RFP filter (top panel) and cold light source (bottom panel). **D)** Percentage of males in the progeny of males carrying the Cas9.1e/Cas9.2a (1e2a), Cas9.1e/Cas9.2c (1e2c) and Cas9.2a/Cas9.2c (2a2c) transgenes crossed to wildtype females, compared to a wildtype cross (WT). The numbers indicate the total number of individuals scored.

We next tested the effects of combinations of transgenes that each individually were able to bias the sex ratio by either targeting the same or both X-linked repeat clusters. We crossed individuals of transgenic strains Cas9.1e, Cas9.2a and Cas9.2c and identified trans-hemizygous individuals using the intensity of polyubiquitin DsRed expression (Figure 3C) as an indicator. We subsequently crossed individual trans-hemizygous males to 10 wild type females and determined the sex ratio of their progeny in 10 replicate crosses for each combination (Figure 3D). We found no substantial boost to the average male bias in the progeny of males carrying the Cas9.1e/Cas9.2c or Cas9.2a/Cas9.2c insertions over and above what had been observed with single doses of the more effective of the two transgenes. This suggests that even an additional dose of Cas9 was not enough to significantly increase the male bias in these two circumstances. In contrast, males carrying the Cas9.1e/Cas9.2a combination showed an average sex ratio of ~80%, suggesting that particular combinations of gRNAs can indeed boost the sex bias substantially.

### Analysis of the target repeat structures

We used a recent re-assembly of the medfly genome built with PacBio and Hi-C data (EgII_Ccap3.2.1)^27^ to characterize the possible origin and structure of the sequences selected by redkmer and in particular of the experimentally targeted kmers. While the assembly remains somewhat fragmented (N50 ~77.38 Mb) we reasoned that it could possibly help to understand the nature and function of the sequences targeted.

We first evaluated chromosomal origin for each of the scaffolds of the EgII_Ccap3.2.1 assembly by calculating the chromosome quotient (CQ) for all 18,520 genes in the annotated EgII_Ccap3.2.1 geneset and then assigned CQ to each of the 2,712 contigs based on the average CQ of genes it contains (Figure 4A, Supplementary Table S4). This analysis revealed that the X-chromosome constitutes the bulk of scaffold 3, but that this scaffold also contains Y-chromosome derived sequences (Figure 4A). We then used BLASTN to locate the redkmer candidate X-kmers, including the top twenty-five kmers and the set of experimentally validated kmers, within the genome assembly. In accordance with expectation, the candidate X-kmers mapped predominantly to scaffold 3 (Figure 4A and Supplementary Figure 2), demonstrating the specificity of X-chromosome targeting by redkmer and our downstream selection. Both experimentally validated Cas9 kmers mapped exclusively to scaffold 3 with 66 and 50 hits for Cas9.1 and Cas9.2, respectively. Cas12a.2 was also specific to scaffold 3 but with only 8 target site hits, while Cas12a.1 also had one hit on scaffold 2 and 15 of scaffold 3 (Supplementary Figure 2). Interestingly, the best match in the assembly for kmer Cas9.1 showed a single-base A to G mismatch in 56 of the 66 target sites, the remaining had additional mismatches (Figure 4B), which may reflect strain differences.

**Figure 4.**
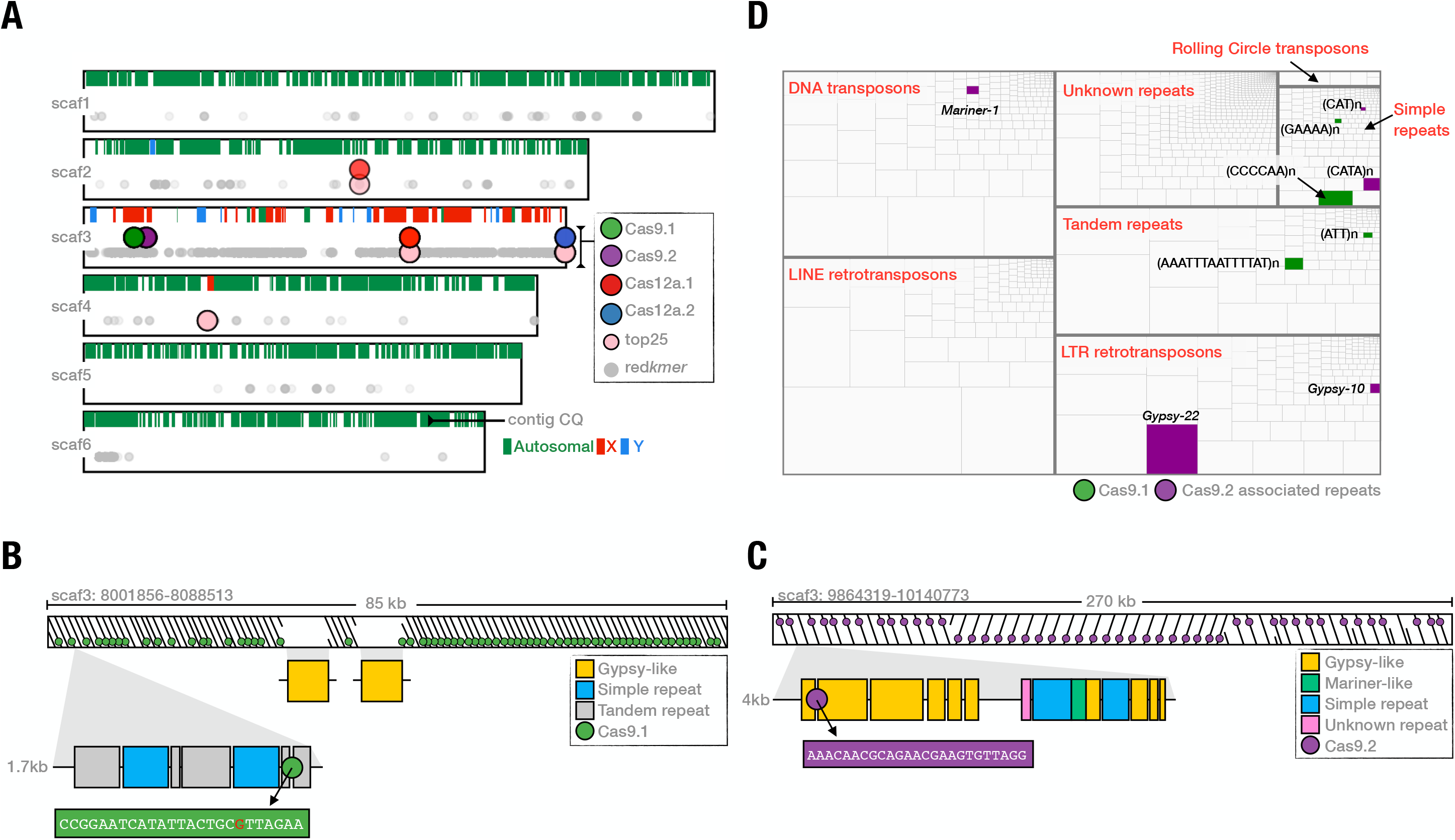
Cas9 kmers target X-chromosome specific repeats. **A)** Distribution of selected kmers in the EgII_Ccap3.2.1 genome assembly. For each of the top six longest scaffolds in the EgII_Ccap3.2.1 genome assembly contig CQ is shown and the position of the kmer hits from either all redkmer output (grey), the top25 selected kmers (pink) and for the four experimentally verified sgRNAs (Cas9.1 in green; Cas9.2 in purple; Cas12a.1 in red; Cas12a.2 in blue). **B)** Organization of the Cas9.1 targeted repeat region. **C**) Organization of the Cas9.2 targeted repeat region. **D)**The medfly genomic repeatome and the distribution of repeats associated with the ~85kb and ~270kb regions targeted by Cas9.1 and Cas9.2, respectively.

To understand the possible origin and function of the sequences we targeted, we next used RepeatModeler^28^ to annotate and catalogue the resident repeats in the EgII_Ccap3.2.1 assembly, since the kmer hits on scaffold 3 did not overlap with any known genome annotations. A library of 2,037 consensus repeats including transposable elements, tandem and low-complexity repeats was mapped on the assembly using RepeatMasker and regions surrounding the Cas9 kmer targets on scaffold 3 were manually annotated. Cas9.1 kmer hits were distributed in a region spanning ~85kb of scaffold 3 (8,001,979-8,087,013), and Cas9.2 kmer hits spanned a region of ~270kb (9,864,418-10,124,412). The Cas9.1 target sequence was embedded in an approximately 1.7 kb long repeat unit, which itself was composed of distinct simple and tandem repeats that were repeated and arranged head-to-tail throughout the array, with two interruptions from a Gypsy-like element (Figure 4B). The Cas9.2 target sequence instead was derived from a larger 4kb repeat unit that is repeated 50 times in a head to tail fashion with a large repeat inversion in the middle (Figure 4C). We found that the 4kb repeat unit is mostly composed of fragmented retrotransposons and simple repeat elements and the Cas9.2 target sequence was located in an incomplete Gypsy-like element. To understand the representation of the targeted repeats within the entire landscape of medfly repeatome, we calculated the relative abundance of each of the consensus sequences of our custom repeat library using RepeatMasker and then highlighted those derived from the 85kb and 270kb regions of the Cas9.1 and Cas9.2 target kmer, respectively (Figure 4D). We found no evidence of active transcription occurring at these two repeat clusters (data not shown). We concluded from these results that X-chromosome shredding sex ratio distorters can be engineered by targeting resident repeats of the X-chromosome and these repeats do not have to be highly abundant, conserved or functional.

## Discussion

We were able to achieve nearly optimal rates of Cas9 activity in the male germline against the *white eye* gene using the *C. capitata* beta2 tubulin promoter and the CcU6.1 promoter we have isolated. The establishment of an efficient CRISPR/Cas9 toolset for transgenesis opens a number of possibilities from the generation of strains for achieving genetic sterility to the development of suppressive gene drives. Activity against *white eye* was lower for strain Cas12a.w, but Cas12a appears a viable genome editing system in the medfly offering a complementary set of targetable sites. No activity against white could be detected for the multiplexed strain Cas12a.m although it shared the same gRNA targeting *white eye* with strain Cas12a.w. This either suggests a strong position effect affecting Cas12a or gRNA expression levels between these two strains or, as previously demonstrated, a loss in efficiency of expression and cleavage when gRNA arrays are utilized^29^. It also does not allow us to conclude on the suitability of the Cas12a.1 and Cas12a.2 target repeats for X-shredding and the former kmer, predicted by redkmer to be an order of magnitude more abundant than both selected Cas9 kmers (Figure 1B), remains an attractive target to be evaluated in the future.

Both Cas9 gRNAs we tested in the medfly gave rise to a significant sex bias, which improves upon previous results in *Drosophila* where only one out eight gRNAs designed for X-shredding were found to be effective^8^. The consistency of the relative strength of the observed bias within the set of Cas9.1 and Cas9.2 strains suggested that the nature of the target sequence was more important in determining the effect on the sex ratio than possible expression differences between different insertion sites of the transgene. This result is in stark contrast with results in *Anopheles* where different integrations generated widely varying levels of expression and distortion^6,7^ and is possibly explained by our use of insulator sequences flanking Cas9. It also suggests that constructs lacking such insulators could be used to select for more active X-shredders.

The target sequences of the two previously studied successful X-shredding systems fell within the coding sequence of the X-linked *Muc14a* gene and within the X-linked rDNA gene cluster, in *Drosophila melanogaster* and *Anopheles gambiae*, respectively. For the present set of experiments in the medfly, our bioinformatic pipeline, making no assumption about the nature or function of the most abundant and X-specific repeats, identified and selected targets that are associated with clusters of X-linked repeats. These repeats, featuring and average distance of approximately 1.7 and 4 kb between the Cas9.1 and Cas9.2 targets respectively, were found to consist of both simple repeat sequences and fragments of retrotransposons. They are unlikely to be functional and presumably constitute heterochromatic regions of the X chromosome. This suggests that sex distorters targeting repeats could be engineered with relative ease in species with an abundance of X-specific heterochromatin, likely a common occurrence in many insects. The fact that the mere presence of a sex chromosome specific repeat sequence constitutes a sufficient substrate for the operation of a highly-efficient distortion trait also supports the notion that the evolution of both X and Y specific sequence heterochromatin could to a large extent drive and in turn be driven by the evolutionary dynamics of sexually antagonistic selfish genes^30^. A downside of selecting such “junk” DNA targets may be the low level of functional conservation which would facilitate the selection for resistance, although the actual mechanism by which X-shredding eliminates X-bearing gametes remains unknown and hence also the pathways via which resistance is likely to arise. Several caveats apply to the post hoc analysis we performed based on the genome assembly and the conclusions we reached above. As mentioned, the actual structure of these repeat clusters may differ in size and composition from what is predicted by the assembly and possibly also between strains and even individuals. Furthermore, the EgII_Ccap3.2.1^27^ assembly was generated from individuals of medfly laboratory strain (EgyptII) which differs from our experimental strain (Benakeion) and the strain used for redkmer target site selection (FAM18). Our abundance estimates and predictions on chromosome specificity and/or size of the target locus may in fact differ between these strains. This is exemplified by Cas9.1 kmer where we observed strain-specific differences between the kmer targets predicted by redkmer and the genome assembly. While the single nucleotide mismatch 16 nucleotides away from the PAM was not predicted to abolish Cas9 activity, the weaker sex distortion effect we observed for Cas9.1 may be partially explained by this discrepancy. The analysis also underlined the importance of a stringent downselection of putative X-specific targets. Both repeat clusters we analysed provided a large set of alternative targetable sites many of which are more abundant than the Cas9 kmers we had selected. A subset of these possible targets also appears to be X-specific i.e. abundant on Scaffold 3 yet absent from the remaining Scaffolds in the assembly (Supplementary Figure 3A,B). Redkmer however excluded such targets due to their lack of X-specificity determined on the basis of long read and short read quotients. It further proves the utility of dedicated tools such as redkmer for the bioinformatic identification of chromosome-specific sequence families e.g. CRISPR target sequences to induce X-shredding for species of medical or agricultural importance for which high-quality genomes are not available.

We demonstrated synergy when a combination of gRNAs targeting different repeats were used, resulting in male bias levels that are closer to what would be required for application. Combining two different gRNAs resulted in higher levels of distortion than any single gRNA or a double dose of a well performing gRNA and Cas9 in homozygous males (Figure 3D). The latter experiment was of particular interest as it has not previously been possible in *An. gambiae* or *D. melanogaster*, where for each organism only a single functional X-shredding target is currently available. This result suggests that further improvements in the CRISPR toolkit for the efficient expression of multiple gRNAs could allow the generation of X-shredders simultaneously targeting a class of less-repetitive X-linked sequences. In *Drosophila* the combination of X-shredding and X-poisoning systems has also been demonstrated and could help to strengthen the robustness of such traits against resistance or escape^8^.

Our findings suggest that an autosomal and hence self-limiting medfly sex distorter strain with performance and fitness characteristics suitable for inundative genetic control in the field could now be developed. Such a strain could conceivably combine the existing genetic sexing system and mass rearing technology of medfly with an inducible or repressible sex distortion system and could decrease the cost of area wide control of the medfly. Finally, the flexibility of the genetic architecture controlling medfly sex determination where both fertile XY females and XX males have been generated^12^, suggests that an attempt to generate a driving Y-chromosome would also be fruitful endeavour in this species. This would not only provide, for the first time, a well-defined system to study Y drive, which is currently lacking altogether, but also a powerful inoculative method of control of this agricultural pest.

## Materials and Methods

### X-shredding target site selection

To select our Cas9 and Cas12a X-chromosome target sequences we ran redkmer^21^ using medfly male and female Illumina whole genome sequencing (WGS) data and canu-corrected^31^ PacBio whole genome sequencing reads from males with the following parameters: TRIMM5=5, TRIMM3=5, pac_length=2000, pac_length_max=100000, LSum=100, XSI=0.995, kmernoise=5. The mitochondrial genome was downloaded from NCBI (NC_000857.1) and used to exclude mitochondrial derived reads. The initial output of redkmer was a set of kmers that were analysed for Cas9 and Cas12a targeting using Flashfry^24^ using as a background genome the PacBio reads deriving from the redkmer autosomal and Y-derived PacBio bins. Candidate X-kmers were then filtered based on their abundance in the Illumina (sum) and PacBio (hits_sum) WGS libraries (Figure 4B). For each candidate X-kmer we also calculated its maximum occurrence in a single PacBio read (max.pt – frequency) and the total number of unique PacBio reads that contain it (Unique_bin_Occurence, Figure 4D). For each of these four parameters (sum, hits_sum, max.pt and Unique_bin_Occurence), kmers occurring in the top 0.05% percentile among the redkmer candidate output kmers were identified and these criteria were then used to select the top 25 kmers with Cas9 or Cas12a targeting potential and excluding those with high off-targets. Scripts used for the filtering filtering are available on the https://github.com/genome-traffic/medflyXpaper.

### Isolation of the *CcU6* promoter

The *D. melanogaster* U6snRNA sequence (GenBank accession no. NR002083) was used to perform a BLAST search of the *Ceratitis capitata* genome (taxid:7213). A sequence 450 bp upstream of the region coding for a predicted U6 spliceosomal RNA in *C. capitata* (LOC111591613, ncRNA, NCBI Reference Sequence: XR_002750719.1) was then selected as the *C. capitata* U6 promoter (*CcU6*).

### Cas9 and Cas12a constructs

The p1260 plasmid (kindly provided by Francesca Scolari, University of Pavia, Italy) with piggyBac recombination sequences and pUb-DsRed as described in Scolari *et al.* (2008)^25^ was used as the backbone of PiggyBac_Cas9 and PiggyBac_Cas12a constructs. For this study we used the following synthetic plasmids and primers by Eurofins genomics: CcU6-gRNA-Cas9; CcU6-gRNA-Cas12a.w; CcU6-gRNA-Cas12a.m. To generate the Cas9 constructs, three DNA fragments were amplified: 1) a 1250 bp fragment including the attP site and Ccβ2-tubulin 5’ regulatory region was amplified from p1260 with primers *β2-promoterF* and *β2-promoterR*; 2) the human codon-optimized Cas9 coding sequence including two nuclear localization signals (SV40 NLS at the 5’ and nucleoplasmin NLS at the 3’) was amplified from hCas9 (Addgene #41815; http://n2t.net/addgene:41815) using primers *F_Cas9* and *R_Cas9*; 3) the 641 bp 3’-UTR of the β2-tubulin gene was isolated from genomic DNA (extracted from a pool of five adult *C. capitata* males using the Holmes-Bonner protocol) using primers *F_3UTR-β2* and *R_3UTR-β2*. The PCR fragment was amplified after purification with primers *F1_3UTR-β2* and *R1_3UTR-β2-SacII*, the latter containing the *SacII* restriction site. These three PCR products were assembled into an *AscI*-linearized p1260 plasmid to create the PiggyBac_Cas9 construct. The CcU6-gRNA-Cas9 plasmid containing the pair of *AarI* restriction sites used for golden gate cloning of the specific gRNA served as the PCR template for the cloning of the individual gRNA expression plasmids using primers containing *SacII* site, *SacII-CcU6-F* and *SacII-CcU6R* and the obtained amplicon was cut with *SacII* and then ligated to *SacII*-linearized plasmid PiggyBac_Cas9. All gRNAs were cloned in *AarI*-linearized plasmid PiggyBac_Cas9 creating the three vectors PiggyBac_Cas9.w, PiggyBac_Cas9.1 and PiggyBac_Cas9.2. The PiggyBac_Cas12a plasmid was generated through three steps. In the first step, the *HindIII*- linearized pUK21 plasmid (Addgene #49788; https://www.addgene.org/49788) was used as an intermediate backbone to clone: 1) the Ccβ2-tubulin-promoter (from p1260) amplified with the forward primer designed with restriction site *MluI*, *1a-Bp-MluI-F*, and the reverse primer *1a-Bp-R*; and 2), 3’UTR β2-tubulin terminator with the forward primer containing the protospacer-*BsaI* site, *BsaI-Bt-F*, and the reverse primer containing the *SacII* site, *Bt-SacII-R*. The two PCR fragments were re-amplified with the following primers with the all characteristics necessary for assembly: *Gb1-pUk21-B2p-F*, *Gb1-pUk21-B2p-R* and *Gb2-BpSpacerBT-F*, *Gb2-BpSpacerBT-R*, respectively. The final products were assembled in pUK-Ccβ2. In the second step, Cas12a (Addgene #69988; http://www.addgene.org/69988/) was amplified with the primers *Cas12a-BsaI-F* and *Cas12a-BsaI-R,* and was assembled through golden gate cloning in the *BsaI*-linerarized plasmid pUK-Ccβ2, generating the plasmid pUKCas12a. In the last step, the pUKCas12a plasmid was cut with the enzyme *MluI* and the digested fragment ligated to the *AscI*-linearized p1260. The CcU6-gRNA-Cas12a.w; CcU6g-RNA-Cas12a.m plasmids, provided with the specific gRNA sequence, were amplified with primers *CcU6-Cas12a-F* and *CcU6-Cas12a-R*. The PCR products were assembled in the final PiggyBac_Cas12a.w and PiggyBac_Cas12.a.m constructs, respectively. The annotated GenBank files for all plasmid vectors are provided in Supplementary File 1, for primers see Supplementary Table S2.

### *Ceratitis capitata* germline transformation

Germline transformation was performed by microinjection of piggyBac constructs (500 ng/μl) together with the ^*i*^*hyPBase*^32^ transposase helper plasmid (300 ng/μl) into the wild-type embryos as described in Meccariello *et al.* (2019)^12^. Hatched larvae were transferred to Petri dishes containing larval food (Meccariello et al 2017)^20^. Surviving G_0_ individuals were crossed to wild type flies, and positive transformants were identified under fluorescence microscopy for the expression of the PUb-DsRed among the G_1_ progeny. Transgenic lines originated from a single integration event were selected using inverse PCR following the protocol as described in Scolari *et al.* (2008)^25^ using the primers from Galizi *et al.* (2014)^6^(Supplementary Table S3). In each generation, transgenic females were backcrossed to *Benakeion* wild-type males. Flies were screened using the fluorescence stereomicroscope MVX-ZB10 Olympus with the filter for RFP (exciter filter 530-560 nm, dichroic beam splitter 570 nm, barrier filter 585-670 nm).

### Medfly rearing

The *C. capitata* transgenic target line, the wild-type strain *Benakeion,* was kindly provided by Giuseppe Saccone’s Lab, University of Naples “Federico II”, and *Benakeion white eye* mutant strain (w2Δ) was provided by the FAO/IAEA Agriculture and Biotechnology Laboratory, Seibersdorf, Austria. The strains were maintained in standard laboratory conditions at 26 °C, 65% relative humidity and 12:12 h light–dark regimen. The adult flies were fed yeast/sucrose powder (1:2).

### Cas9.w and Cas12a activity assays against *white eye*

A transgenic male *we^+^* homozygote was individually crossed to five homozygous w2Δ mutant females and, as a control, a transgenic female *we^+^* homozygote was individually crossed to five homozygous w2Δ mutant males; a wild-type male was individually crossed to five homozygous w2Δ mutant females; a wild-type female was individually crossed to five homozygous w2Δ mutant males and the number of white eye phenotype adults in the progeny was counted. The crosses of the Cas12a transgenic lines were set up at 34 °C, 65% relative humidity and 12:12 h light–dark regimen. For the molecular analysis of CRISPR/Cas9-induced mutations in *white eye* gene genomic DNA was extracted from individual flies in 200 μl Holmes Bonner buffer, with minor modifications, according to the protocol of Holmes and Bonner (1973)^33^. The resulting DNA was used as template to amplify the region encompassing the target sites, using the following primers: Cas9_W-F and Cas9_W-R. The PCR products were purified with Monarch® PCR & DNA Cleanup Kit (New England Biolabs) and subcloned using StrataClone PCR cloning Kit (Agilent Technologies) and sequenced.

### Determination of the sex ratios and egg-to-adult survival assay

To assay the adult sex ratio, 20 transgenic males were crossed to 30 wild-type females and, as a control, 30 transgenic females were crossed to 20 wild-type males. Each transgenic line was assayed for at least four consecutive generations. Eggs were collected three times, with an interval of two to three days and reared to adulthood. The total number of adult male and female flies from each cross were counted. The survival test was performed, 20 transgenic males were crossed to 30 wild-type female and as a control 20 wild-type males were crossed to 30 wild-type females. The number of eggs laid as well as the number of adults hatching were counted. Each transgenic line was assayed for one generation. Three different crosses were performed to generate individuals with double transgene expression for analysis of the effect of the copy number of the transgene: 1) 10 Cas9.1e males and 20 Cas9.2a females; 2) 10 Cas9.2c males and 20 Cas9.1e females; 3) 10 Cas9.2a males and 20 Cas9.2c females. From each cross 10 males were selected and separated from single insertion of the transgene according to the intensity of DsRed fluorescence and were individually crossed to 10 wild-type females. Egg collection and the determination if the adult sex ratio was in the progeny was performed as previously described. To analyze the sex ratios observed we used a generalized linear model in which the sex-ratios were modelled as a function of the sex of the transgenic parent and the experimental generation using a binomial error distribution. (R scripts and input data are available at https://github.com/genome-traffic/medflyXpaper)

### Reagents

All amplification steps were performed using Phusion High-Fidelity DNA Polymerase (New England Biolabs). The cloning was performed with the NEBuilder Hi Fi DNA Assembly kit (New England Biolabs). The enzymes mentioned were purchased from New England Biolabs: *AscI* (#R0558), *MluI* (#R3198), *BsaI* (#R3733), *SacII* (#R0157). The *AarI* (ER1581) restriction enzyme type II was purchased from ThermoFisher scientific. All inserts were verified by sequencing (Genewiz).

### Origin of the targeted X-chromosome sequences in the EgII_Ccap3.2.1 medfly genome assembly

To identify sequences of the medfly X-chromosome within the EgII_Ccap3.2.1 assembly we calculated the chromosome quotient for each of the 18,520 augustus-predicted genes of the assembly using the EgyptII male and female WGS Illumina data^27^. WGS data from males and females were mapped using bowtie^34^ following the standard CQ parameters (reporting all alignments (-a) and allowing no mismatches (-v 0)) and CQ was calculated as the ratio of female over male (library-size normalized) reads. Contig CQ was then calculated as the median CQ of genes contained in the 2,712 contigs, excluding all contigs containing less than 10 predicted genes. Doing so resulted in median CQ values for the 465 largest contigs representing 15,809 genes (Supplementary Table S4). To generate a custom repeats library for the medfly, we used the assembly of the medfly genome and ran RepeatModeler (v2.0.1-0)^27^, using RECON (v1.08)^35^, RepeatScout (v1.0.6)^36^, and TRF (v4.09)^37^, with default parameters. The consensus classes were compiled in a set of 2,037 sequences belonging to families of transposable elements (TEs), tandem and other low-complexity repeats. Out of the medfly 2,037 reference REs library, 1,312 consensus classes were annotated and classified hierarchically as TEs, Tandem Repeats, and Simple Repeats sequences (https://www.dfam.org/classification/tree and Supplementary Table S5). kmers were mapped to the Ccap3.2 assembly using BLASTN (-word_size 5, -max_target_seqs 10,000 and -max_hsps 10,000). To dissect the structure of the Cas9 targeted repeats in the ~85kb and ~270kb arrays of scaffold 3, we extracted the full sequences 100 bp upstream of the first kmer until the next kmer using the Seqtk toolkit (https://github.com/lh3/seqtk) and then mapped it back to the array using Yass^38^. To annotate the higher order repeat landscape of the ~85kb and ~270kb arrays of scaffold 3, we used RepeatMasker^39^ with default parameters and our previously generated repeats database with the -lib option to generate a gff file. This data was used to map the repeats within the targeted arrays. Additionally, we BLASTed the two Cas9 kmers directly against the RepeatModeler library (-word_size 5, 90% identity, and 50% of coverage) and found a unique hit from the Cas9.1 kmer with a tandem repeat sequence and for the Cas9.2 kmer a significant alignment with an incomplete gypsy-like element. We also ran a BLASTN search (cut-off at 80% identity and e-value of 1e-10) using each of the ~85kb and ~270kb arrays against a merged repeats library that included both the medfly RepeatModeler library and the *Drosophila melanogaster* transposon database (https://github.com/bergmanlab/transposons/tree/master/current). Both approaches, RepeatModeler and BLASTN gave similar results. To generate the genome-wide medfly repeats landscape plot shown in Figure 4D, we used RepeatMasker output to calculate the relative percentage of the total genome assembly masked by each repeat in the library. All repeats were assigned Dfam hierarchical classification nomenclature (full name, sub name, section, type, subtype, section, subsection, and name) to cluster the information into boundary boxes. The percentage of each element associated with the kmers was plotted in R using the treemap package and ggplot (supplemental R scripts and input data are available at https://github.com/genome-traffic/medflyXpaper). To evaluate alternative kmers originating from the targeted regions – the 85 and 270kb regions of scaffold 3 for Cas9.1 and Cas9.2, respectively, we generated 25bp kmers from the genome assembly of these regions using jellyfish count (-C, −c 3 and −s 1000000000 parameters), “dump” (-c) and “histo” (ref). The resulting kmers were then mapped by BLASTN either back to the entire assembly, scaffold 3 alone, or just the region originally used for generating the kmers by jellyfish. Number of hits against each library was extracted from the output and the specificity to either scaffold 3 was calculated.

## Acknowledgements

This study was funded by the BBSRC under the research grant BB/P000843/1 to N.W and by the United States – Israel Binational Agricultural Research and Development Fund (Research Grant No. IS-5180-19) to P.A.P. P.A.P. was also supported by the Italian Ministry Education, University and Research (MIUR—D.M. no. 79 04.02.2014). The funders had no role in study design, data collection and analysis, decision to publish, or preparation of the manuscript.

---

**Supplementary Figure 1. Predicted transgene integration sites within the EgII_Ccap3.2.1 assembly.** Strains are coloured by transgenic construct. Also shown is the location the white-eye gene on scaffold 5.

**Supplementary Figure 2. BLASTN hits of selected kmers to the Ccap3.2 assembly.** Hits are filtered for each of the top six largest scaffolds.

**Supplementary Figure 3. Specificity and abundance of kmers originating in the target regions.** Number of BLASTN hits within scaffold 3 (X-axis) and the entire genome assembly (Y-axis) of kmers originating in the 85 (A) and 270 kb (B) regions. Scaffold 3 specificity is indictated (100% red to 0% blue). Shown in black are those kmers that are specific to the region targeted (hits only in this region within scaffold 3). In green are the two experimentally verified kmers, Cas9.1 and Cas9.2.

