## Supplementary material for "Engineered sex distortion in the global agricultural pest *Ceratitis capitata*": Supplementary Table S3.docx

| Transformation construct | Strain | Scaffold_id | Annotation_  CCAP3.2_CANU | Insertion site |
| --- | --- | --- | --- | --- |
| PiggyBac_Cas9.w | Cas9.w | 2 | 11 598 327 | ACTTTGTCGGTTAA-**PB**-TTAAAATTATGAGC |
| PiggyBac_Cas9.1 | Cas9.1a | N/A |  | - |
|  | Cas9.1b | 5 | 31 827 778 | GAAAATGAATTTAA-**PB**-TTAAATAAAAAACT |
|  | Cas9.1c | 1 | 2 780 832 | GCAAATGAATTTAA**-PB**-TTAAGAGAGAACAA |
|  | Cas9.1d | 6 | 45 939 184 | TGAAACTTGGTTAA-**PB**-TTAAAGCTAAAATC |
|  | Cas9.1e | 4 | 25 561 321 | TAATTAGGTATTAA**-PB-**TTAAAGAGAAAAGA |
|  | Cas9.1f | 4 | 58 949 141 | AAAAAAGTCATTAA-**PB**-TTAAAAGATCAATA |
|  | Cas9.1g | N/A |  | - |
| PiggyBac_Cas9.2 | Cas9.2a | 2 | 78 745 789 | CAATTACATTTTAA-**PB**-TTAACAAGCTCGTT |
|  | Cas9.2b | 2 | 16 554 608 | TATATTTTTTTTAA-**PB**-TTAAAGAAAATAAA |
|  | Cas9.2c | 2 | 19 290 016 | GGATATCCTATTAA-**PB**-TTAAATTCATTAAG |
|  | Cas9.2d | 2 | 70 687 722 | ATCCATTCTCTTAA-**PB**-TTAATAACTCTTGG |
| PiggyBac_Cas12a.w | Cas12a.w | N/A |  | - |
| PiggyBac_Cas12a.m | Cas12a.m | N/A |  | - |
