## Supplementary figures and images for "Engineered sex distortion in the global agricultural pest *Ceratitis capitata*"

### Supplementary Figure 1.pdf

#### Supplementary Figure 1

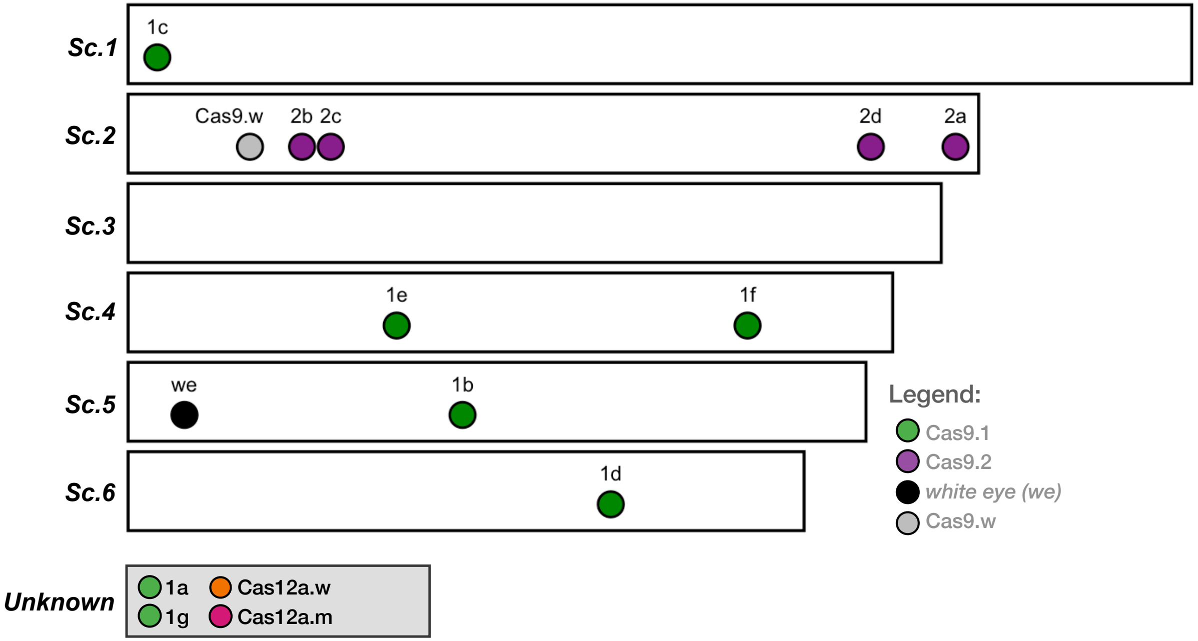

### Supplementary Figure 2.pdf

#### Supplementary Figure 2

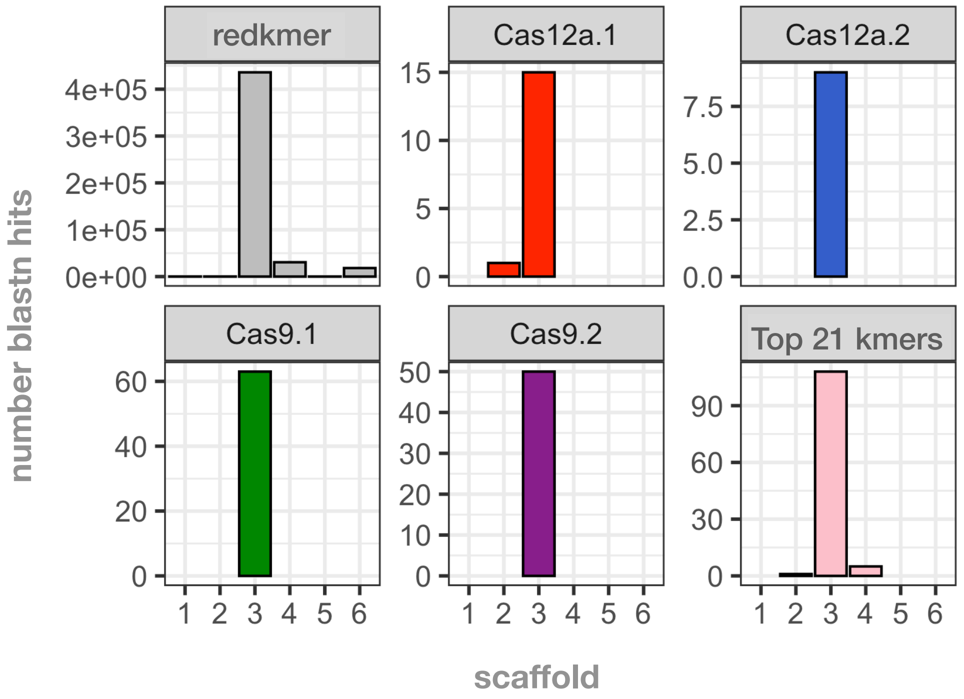

### Supplementary Figure 3.pdf

#### Supplementary Figure 3

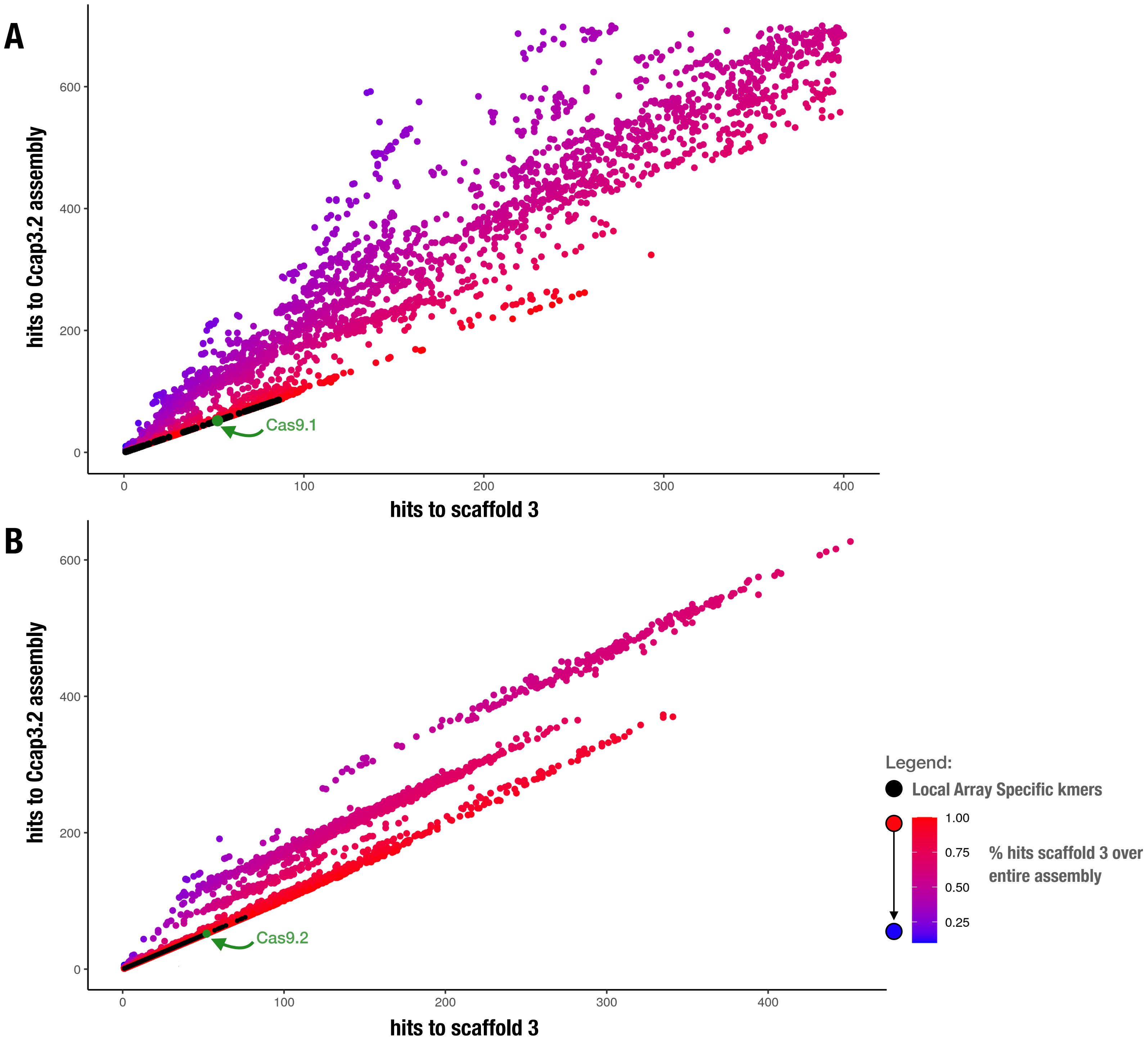
